# NMDA receptor signalling controls R-type calcium channel levels at the neuronal synapse

**DOI:** 10.1101/2020.11.25.399212

**Authors:** Oleg O. Glebov

**Affiliations:** Institute of Neuroregeneration and Neurorehabilitation, Qingdao University, Qingdao 266071, Shandong, China; Department of Old Age Psychiatry, The Institute of Psychiatry, Psychology & Neuroscience, King’s College London, De Crespigny Park, Denmark Hill, London SE5 8AF, UK

## Abstract

Regulation of extracellular Ca++ influx by neuronal activity is a key mechanism underlying synaptic plasticity. At the neuronal synapse, activity-dependent Ca++ entry involves NMDA-type glutamate receptors (NMDARs) and voltage-gated calcium channels (VGCCs); the relationship between NMDARs and VGCCs, however, is poorly understood. Here, I report that neuronal activity specifically regulates synaptic levels of R-type VGCCs through synaptic NMDAR signalling and protein translation. This finding reveals a link between two key neuronal signalling pathways, suggesting a feedback mode for regulation of Ca++ signalling at the synapse.

## Introduction

The calcium (Ca++) ions play a central role in the regulation of the synaptic function in the central nervous system (CNS). Upon arrival of the action potential to the synapse, entry of extracellular Ca++ through voltage-gated calcium channels (VGCCs) results in docking of the synaptic vesicles to the presynaptic active zone and the release of the neurotransmitter into the synaptic cleft, resulting in synaptic transmission [1,2]. On the other side of the synapse, depolarization of the membrane by opening of the neurotransmitter receptors opens up VGCCs as well as NMDA-type glutamate receptors (NMDARs), which enable Ca++ to enter the dendritic spine. Ca++ then triggers a complex cascade of signalling pathways that can relay information as far as the cell nucleus, regulating multiple aspects of cell biology such as gene expression, membrane trafficking and protein turnover [3].

NMDARs and VGCCs are the two main sources of Ca++ entry into the synapse. VGCCs are complex proteins possessing multiple transmembrane domains, which open up to allow passage of Ca++ once the cell membrane has depolarized beyond a certain level [1,4]. On the other hand, NMDA receptors sense both depolarization of the membrane and release of glutamate, which enables them to open and allow for entry of various cations, including Ca++ [5,6]. The properties of these channels are therefore quite different. Furthermore, localization of these channels at the synapse also displays notable differences: while NMDARs are specifically located at the postsynaptic dendritic spine, VGCC can be found on both sides of the synapse.

Fast high-voltage VGCCs containing a Cav2 subunit (Cav2-VGCCs) are of particular importance to synaptic Ca++ signalling, as they specifically localize to the synapse. Amongst the three known types of Cav2-VGCCs, P/Q-type (Cav2.1) and N-type (Cav2.2) VGCCs are found on the presynaptic side [7–10], while R-type (Cav2.3) channels operate on both sides of the synapse [11–13]. Despite the well-established roles for both NMDARs and Cav2-VGCCs, the relationship between their signalling pathways remains poorly understood. Local NMDAR and VGCC signalling can be immediately coupled through the short-term biophysical mechanisms [14,15]. On a timescale of days, chronic levels of neuronal activity engage the mechanisms of homeostatic plasticity to regulate many aspects of synaptic protein composition, including all three types of Cav2-containing VGCCs [10,16]. However, the link between NMDAR activity and synaptic Cav2-VGCCs levels remains unclear, and do the underlying cell biological mechanisms.

## Materials and Methods

### Materials

Cell culture reagents were from Invitrogen (UK). Anisomycin, MK801 and memantine were from Sigma Aldrich (UK). TTX, NBQX, APV, and gabazine were from Tocris (UK).

Below is the list of the antibodies used in this study:

| Antigen | Conjugation | Species | Manufacturer | Cat. No. |
| --- | --- | --- | --- | --- |
| Cav2.3 |  | Rabbit | Synaptic Systems | 135302 |
| PSD95 |  | Mouse | Abcam | 13552 |
| Bassoon |  | Mouse | Abcam | 82958 |
| Cav2.1 |  | Rabbit | Synaptic Systems | 152103 |
| Cav2.2 |  | Rabbit | Alomone | ACC-002 |
| Rabbit IgG | Cy5 | Donkey | Jackson Immuno | 711-175-152 |
| Mouse IgG | AlexaFluor488 | Goat | Invitrogen | A-11001 |

### Neuronal culture

Dissociated hippocampal neuronal cultures were isolated from rat embryos at day 18 of gestation and grown according to the Brewer method. Culture medium was Neurobasal with GlutaMax and B-27. No anti-mitotic agents or antibiotics were used during culture. All experiments were carried out at 16-21 days in culture. Cells were cultured on 13 mm poly-L-lysine coated round glass coverslips with 1.5 thickness placed into 35 mm Petri dishes, 4 coverslips per dish. To minimise variability in culture conditions, each experiment was carried out using the cells from the same dissection and cultured within the same Petri dish. All experimental protocols were performed following the guidelines of the local Research Ethics Committee.

### Immunostaining and confocal microscopy

All steps were performed at room temperature (RT). After treatment, coverslips were fixed with 2% (when probing for Psd95) or otherwise 4% para-formaldehyde dissolved in phosphate buffered saline (PBS). Fixation was carried out for 15−20min and was followed by the permeabilization/blocking step. Permeabilization/blocking was carried out in 0.2% Triton-X100 dissolved in PBS supplemented with 5% horse serum, for 10 min. All subsequent incubations were carried out in 0.2% Triton-X100 dissolved in PBS supplemented with 5% horse serum. For primary labelling, coverslips were incubated with appropriate primary antibodies for 45-90 min, then washed 4 times in PBS. For secondary labelling, coverslips were incubated with the appropriate secondary antibodies labelled with AlexaFluor-488 and AlexaFluor-647 at a concentration of 0.3 μg/mL each for 45-90 min. Coverslips were then mounted in Fluoromount-G mounting medium and imaged on a Zeiss LSM710 laser confocal microscope equipped with a standard set of lasers. The imaging system was controlled by ZEN software. Acquisition parameters were as follows: plan-Apochromat 63x/1.4 Oil objective, regions of interest sized 1024×1024 pixels (65.8 nm/pixel), 12-bit, speed 7, averaging setting 2. Excitation laser wavelengths were 488 and 633nm. Bandpass filters were set at 500−550nm and 650−750nm for AlexaFluor-488 and AlexaFluor647 respectively. Pinhole size was kept to 1-2 Airy units. Detector gain settings were optimized to ensure appropriate dynamic range, low background and sufficient signal/noise ratio.

### Image analysis

Image analysis was carried out using the ImageJ software package, version 1.42. Non-synaptic regions with high background fluorescence (e.g. cell bodies) were manually excluded from analysis. For identification of synapses, images were semi-automatically thresholded using the “Moments” setting. Individual synapses were then identified automatically using the “Analyze Particles” command. To avoid rare overlap of multiple synapses, only synapses with areas ranging from 0.1 to 2μm^2^ were included in further analysis. Spatial parameters of the identified synapses were then added to the Region of Interest (ROI) Manager. Individual ROIs were then combined into one compound ROI using the “Combine” and “Add” functions of the ROI Manager interface, whereupon quantification of mean signal intensity in each channel was performed using the “Measure” function. Background subtraction was performed as appropriate.

### Statistical analysis

All the experiments were repeated 3 to 5 times, 5 images per condition. For statistical analysis, Prism 6.0c software package (GraphPad Software) was used. Data distributions were assessed for normality using d’Agostino and Pearson omnibus tests. Student’s t-test and 1-way ANOVA were used for normally distributed datasets to assess statistical significance; for not normally distributed datasets, Mann-Whitney rank test was used. Dunnett’s post-test was used to assess statistical significance of the treatment effects relative to the untreated control samples. Graph plots show mean values and standard error from the mean (SEM).

## Results&Discussion

Blockade of action potential firing by TTX (2uM) for 1 hour caused a significant increase in the Cav2.3 levels within the puncta of a canonical synaptic marker PSD95 (**Fig 1a,b**), although the synaptic level of PSD95 itself was unchanged (**Fig. S1a**). Blockade of inhibitory transmission by a GABA receptor blocker Gabazine (50uM) also increased the synaptic levels of Cav2.3, while moderate depolarization of the neuronal membrane by elevated (15mM) concentration of KCl had no significant effect (**Fig 1a,b**). This indicated that neuronal activity regulated synaptic levels of R-VGCCs.

**Figure 1.**
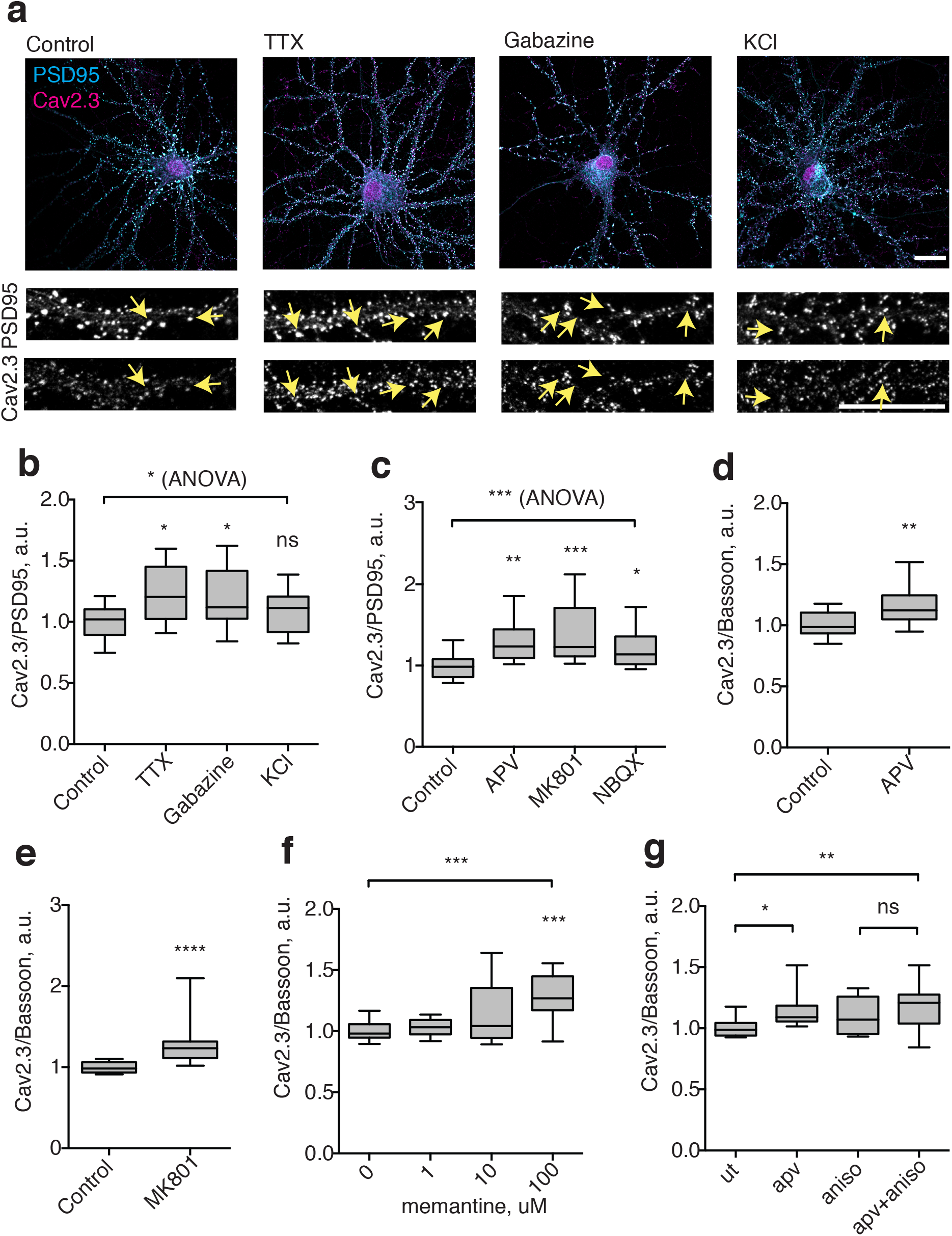
Synaptic NMDAR signalling controls levels of synaptic R-VGCCs. **a**, cells were treated with 2uM TTX, 50uM gabazine or 15mM KCl for 1h at 37C, fixed and immunostained for PSD95 and Cav2.3. Arrows highlight accumulation of Cav2.3 in PSD95-positive puncta corresponding to the synapses. Scale bar, 20um. **b**, quantification of Cav2.3 intensities in PSD-95 positive puncta normalized to PSD95 intensity. ns – not significant, *p<0.05, 1-way ANOVA and Dunnett’s post-test, N=15 fields of view from 3 independent experiments. **c**, Cells were treated with 20uM NBQX, 50uM APV or 20uM MK801 for 1h at 37C. Quantification of Cav2.3 intensities in PSD-95 positive puncta normalized to PSD95 intensity. Ns – not significant, ***p<0.001, **p<0.01, *p<0.05, 1-way ANOVA and Dunnett’s post-test, N=15 fields of view from 3 independent experiments. **d**, Cells were treated with 50uM APV for 1h at 37C. Quantification of Cav2.3 intensities in Bassoon-positive puncta normalized to Bassoon intensity. **p<0.01, Student’s t-test. N=20 fields of view from 4 independent experiments. **e**, cells were treated with 50uM MK801 for 1h at 37C. Quantification of Cav2.3 intensities in Bassoon-positive puncta normalized to Bassoon intensity. ****p<0.0001, Mann-Whitney test. N=20 fields of view from 4 independent experiments. **f**, Cells were treated with varying concentrations of memantine for 1h at 37C. Quantification of Cav2.3 intensities in Bassoon-positive puncta normalized to Bassoon intensity. ***p<0.001, 1-way ANOVA and Student’s t-test. N=15 fields of view from 3 independent experiments. **g**, Cells were treated with 50uM APV and 10uM Anisomycin for 1h at 37C. Quantification of Cav2.3 intensities in Bassoon-positive puncta normalized to Bassoon intensity. **p<0.01, *p<0.05, ns not significant, 1-way ANOVA and Student’s t-test. N=20 fields of view from 4 independent experiments.

To investigate the mechanism coupling neuronal activity and synaptic R-VGCCs, AMPA-type glutamate receptors, which carry out most of glutamatergic synaptic transmission in hippocampus, were blocked using NBQX (20uM). The observed increase in synaptic Cav2.3 plevels resembled that of TTX, suggesting that R-VGCCs leves were reguated by excitatory synaptic transmission (**Fig. 1c**). To further elucidate the signalling mechanism regulating R-VGCCs downstream of synaptic transmission, NMDAR signaling was inibited by a specific blocker APV (50uM). The resulting effect was similar to that of TTX and NBQX, indicating that blockade of NMDARs was sufficient to induce recruitment of R-VGCCs to the synapse. Importantly, treatment with a structurally unrelated NMDAR blocker MK801 (20uM) had the same effect as APV, further confirming involvement of NMDAR signalling in synaptic R-VGCC regulation (**Fig. 1c**).

To further confirm synaptic enrichment of Cav2.3, we quantified the Cav2.3 levels in areas labeleed by a different synaptic marker Bassoon (Bsn). Treatment with APV or MK801 significantly increased Cav2.3 relative to Bsn levels, indicating that the change in the ratio was indeed caused by the specific increase in synaptic levels of Cav2.3 (**Fig. 1d,e**). This conclusion was further supported by an increase in the somatic levels of Cav2.3, suggesting that the levels of R-VGCCs were increased across the cell rather than only at the synapse (Fig). (p<0.05, Mann-Whitney test, **Fig. S1b**).

Other types of Cav-2 containing VGCCs, namely N-VGCCs and P/Q-VGCCs, have been shown to slowly accumulate at the synapse over the course of 24-48h upon blockade of activity, consistent with the timescale of homeostatic plasticity [10,16]. To investigate whether their timescale of their recruitment matched that of R-VGCCs, immunostaining for the pore-forming subunits Cav2.1 and Cav2.2 was performed in cultures treated with APV for 1h (**Fig. S1c,d**). Synaptic levels of both Cav2.1 and Cav2.2 were not significantly increased, suggesting that the accumulation of VGCCs at the synapse triggered by the 1h NMDAR blockade is restricted to R-VGCCs and does not affect N-VGCCs and P/Q-VGCCs (**Fig. S1e,f**).

Besides synaptic NMDARs, activation of extrasynaptic NMDARs has also been implicated in physiologically and clinically important sugnalling pathways, including neurotoxicity. To differentiate between these two modes of NMDAR signalling, I took advantage of the pharmacological profile of the drug memantine, which preferentially inhibits extrasynaptic NMDAR in low concentrations [17]. Treatment with low concetrations of memantine (1-10uM) had no effect on Cav2.3 accumulation, whereas a higher concentration (100uM) significantly increased synaptic Cav2.3 (**Fig. 1f)**. Therefore, it can be concluded that synaptic rather than extrasynaptic NMDARs signalling regulates synaptic R-VGCCs.

A major mechanism for NMDAR-dependent regulation of synaptic composition is through translational control, whereby Ca++ influx through NMDAR activity limits protein elongation [18]. To test for the role of translation, neurons were treated with anisomycin (10uM), a well-established blocker of protein elongation. Treatment with anisomycin abolished the APV-induced increase in synaptic Cav2.3, suggesting that translation was indeed required for the increase of synaptic R-VGCCs triggered by the NMDAR blockade (**Fig. 1g**).

This study reports that neuronal activity rapidly controls levels of R-VGCCs in the synapse through excitatory synaptic transmission, synaptic NMDAR signalling and translation. This regulation is likely to be indirect, given that translation rate of Cav2.3 itself has been shown to be activity-independent [19]. Nevertheless, spatial proximity between NMDARs and R-VGCCs at the postsynaptic compartment [13,15] would suggest the involvement of local processes. In agreement with this notion, the timescale of the observed coupling between NMDARs y and R-VGCCs (within 1h) is considerably faster than the previously reported slow (24-48h) mechanisms of homeostatic plasticity regulating presynaptic Cav2-containing VGCCs [10,16,20]. In contrast, P/Q and N-VGCCs are primarily presynaptic, and the slower timescale of their regulation by neuronal activity likely reflects involvement of other mechanisms, which may operate either presynaptically or across the cell.

Upregulation of synaptic R-VGCCs by both blockade of excitatory NMDAR receptors as well as blockade of inhibitory GABA receptors may seem paradoxical (**Fig. 1b**), given that blocking of inhibitory neurotransmission by GABA receptor antagonists immediately increases neuronal firing [21]. However, this effect is consistent with blockade of GABAergic transmission triggering activity-induced reduction in synaptic NMDAR content [22,23]. Therefore, it may be concluded that three different ways of downregulation of synaptic NMDAR function – due to either blockade of neuronal firing, or homeostatic reduction in synaptic NMDAR levels through increased neuronal firing, or simply by direct pharmacological blockade of synaptic NMDARs – all lead to the same outcome, i.e. upregulation in synaptic R-VGCCs.

Given the spatial constraints dominating local signalling at the synapse, restriction of local Ca++ signalling through the feedback mechanim reported here is likely to be a key factor allowing for co-existence of multiple signalling pathways within the synapse through tuning synaptic Ca++ signalling to neuronal activity [13]. Considering the major role of synaptic Ca++ signalling in neuronal development, plasticity, and pathology, observations reported here will warrant deeper investigation of the relationship between local NMDAR and VGCC signalling. This will be of particular interest in functionally relevant contexts implicating R-type VGCCs, e.g. neuronal development [24], physiologically validated forms of synaptic plasticity, and neuronal pathology [11,25,26].

## Data availability statement

The data that support the findings of this study are available from the corresponding author upon reasonable request.

## Figure legends

**Figure S1.**
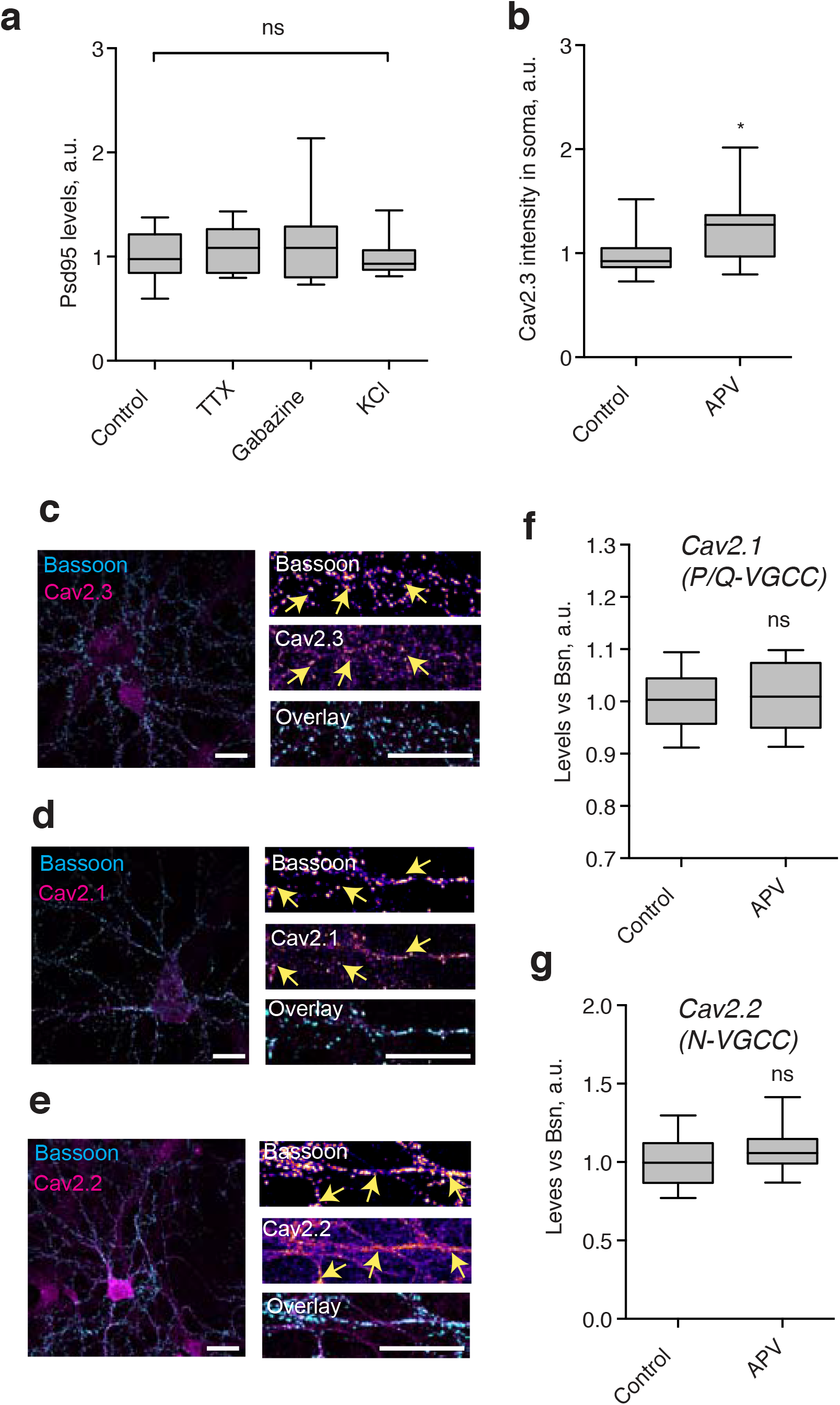
Extended experimental data. **a**, quantification of PSD-95 intensities in PSD-95 positive puncta. ns – not significant, 1-way ANOVA and Dunnett’s post-test, N=15 fields of view from 3 independent experiments. **b**, Accumulation of Cav2.3 labelling in the soma of neurons following APV treatment. *p<0.05, Mann-Whitney test, N=15 cells from 3 independent experiments. **c**, Neurons were immunostained for Cav2.3 and Bassoon. Arrows highight accumulation of Cav2.3 signal in Bassoon-positive puncta. Scale bar, 20um. **d**, ditto for Cav2.2. **e**, ditto for Cav2.2. **f**, Lack of enrichment of Cav2.1 in Bassoon-positive areas following APV treatment. ns - not significant, Mann-Whitney test. N=15 fields of view from 3 independent experiments. **g**, Lack of enrichment of Cav2.2 in Bassoon-positive areas following APV treatment. ns - not significant, Mann-Whitney test. N=20 fields of view from 4 independent experiments.

